# A role for Fibroblast Growth Factor Receptor 1 in the pathogenesis of *Neisseria meningitidis*

**DOI:** 10.1101/283028

**Authors:** Sheyda Azimi, Lee M. Wheldon, Neil J. Oldfield, Dlawer A. A. Ala’Aldeen, Karl G. Wooldridge

## Abstract

*Neisseria meningitidis* remains an important cause of human disease. It is highly adapted to the human host – its only known reservoir. Adaptations to the host environment include many specific interactions with human molecules including iron-binding proteins, components of the innate and adaptive immune systems, and cell surface receptors such as the Epidermal Growth Factor Receptor (EGFR). Interaction of the meningococcus with EGFR has been elucidated in some detail and leads to intracellular signalling and cytoskeletal changes contributing to the pathogenesis of the organism. Here, we show that the meningococcus also recruits Fibroblast Growth Factor Receptor 1 (FGFR1) onto the surface of human blood microvascular epithelial cells (HBMECs). Furthermore, meningococci internalised into these cells recruit the activated form of this receptor, and that expression and activation of FGFR1 is necessary for efficient internalisation of meningococci into HBMECs. We show that *Neisseria meningitidis* interacts specifically with the IIIc isoform of FGFR1.

## Introduction

*Neisseria meningitidis*, or the meningococcus, while normally a harmless commensal of the human oropharynx, can occasionally cause devastating disease including meningitis, sepsis, disseminated intravascular coagulation (DIC) and multiple organ failure (Takada *et al.*, 2016). Penetration of meningococci through the oropharyngeal epithelial mucosa and into the blood is a crucial step in the development of sepsis and other systemic diseases, while penetration of the blood-brain barrier (BBB) is a prerequisite for the development of meningitis (Nassif, 1999, Virji, 2009). Attachment to endothelial cells induces membrane protrusions at the bacterial binding site and leads to the formation of specific protein complexes known as cortical plaques underneath the bacterial colonies (Merz *et al.*, 1999, Eugene *et al.*, 2002). The process by which these steps occur is incompletely understood (Sokolova *et al.*, 2004, Yazdankhah *et al.*, 2004, Hill *et al.*, 2010), but identified meningococcal adhesins include components of the type IV pilus (PilC1, PilC2 and PilQ), the outer membrane proteins Opa, Opc, Factor H-binding protein, PorA, PorB, HrpA, NadA, App and MspA, as well as lipooligosaccharide (LOS) (Merz *et al.*, 2000, Hadi *et al.*, 2001, Turner *et al.*, 2006, Morand *et al.*, 2009). Host cell receptors that have been identified include alpha actinin, integrins, CEACAMS, CD46, Complement receptor 3, GP96 scavenger receptor, laminin, platelet-activating factor, mannose receptor, Transferrin receptor 1, Laminin receptor and Galectin-3 (Merz *et al.*, 2000, Morand *et al.*, 2009, Orihuela *et al.*, 2009, Quattroni *et al.*, 2012, Alqahtani *et al.*, 2014, Khairalla *et al.*, 2015).

Fibroblast Growth Factor Receptors (FGFRs) are transmembrane proteins that belong to the Receptor Tyrosine Kinase (RTK) family of signalling molecules. This family consists of four members that are responsible for recognising all 22 Fibroblast Growth Factor molecules (FGFs) found in humans (Ornitz *et al.*, 2001, Turner *et al.*, 2010, Guillemot *et al.*, 2011). FGFs are involved in cell differentiation, migration and proliferation during early embryogenesis and play an important role in tissue repair, wound healing (Ortega *et al.*, 1998) and tumour angiogenesis in adulthood (Gerwins *et al.*, 2000, Eswarakumar *et al.*, 2005, Presta *et al.*, 2005). Splicing of FGFR transcripts generates a variety of specific isoforms in various types of cells and tissues recognising specific types of FGF molecules (Miki *et al.*, 1992, Groth *et al.*, 2002, Eswarakumar *et al.*, 2005, Turner *et al.*, 2010).

FGFRs are single transmembrane receptors; the extracellular N-terminal region consists of three IgG-like domains that form a ligand-binding domain with an acidic box which interacts with heparin sulphate proteoglycans (HSPGs) and Cell Adhesion Molecules (CAMs). This is followed by a transmembrane region and a C-terminal cytoplasmic region containing 7 specific tyrosine residues (Ornitz *et al.*, 1992, Reiland *et al.*, 1993, Cavallaro *et al.*, 2004, Francavilla *et al.*, 2009). Binding of FGFs to FGFR leads to dimerisation of the receptor and activation of tyrosine autophosphorylation, which in turn activates the receptor, leading to activation of downstream signalling pathways (Mohammadi *et al.*, 1992, Mohammadi *et al.*, 1996, Lundin *et al.*, 2003, Zhang *et al.*, 2006). Previous studies confirmed the importance of FGFR1 signalling and expression in maintaining the integrity and differentiation of endothelial cells forming the microvasculature (Kanda *et al.*, 1996, Gerwins *et al.*, 2000, Murakami *et al.*, 2008).

The possible role of FGFRs in infectious diseases has not been excessively investigated, although FGF2 expression enhance *Chlamydia trachomatis* binding and internalisation into epithelial cells (Kim *et al.*, 2011). *C. trachomatis* facilitates entry by binding directly to FGF2, which results in binding of FGF2-bacteria-heperan sulfate proteoglycan (HSPG) complexes to FGFR and internalisation of the elementary bodies of these bacteria into the epithelial cells. The study also showed higher levels of FGFR substrate 2 (FRS2) activation as a results of FGF2/*C. trachomatis* treatment in Hela cells, and higher level of FGF2 expression via activation of ERK1, 2. More recently, HSPG-associated FGFR1 has been implicated in internalisation of *Rickettsia rickettsii* into cultured human microvascular endothelial cells and inhibition of FGFR1 in a *R. conorii* murine model of endothelial-target spotted fever rickettsiosis reduced the rickettsial burden in infected mice (Sahni *et al.*, 2017).

In a study of role of RTKs, and specifically EGFRs, in meningococcal infection higher levels of FGFR1 activation were observed in endothelial cells in response to infection (Slanina *et al.*, 2014). Here we investigated the possible direct interaction of FGFRs expressed in HBMECs with meningococci and the influence of such an interaction on the ability of *N. meningitidis* to invade these cells.

## Results

### Meningococci recruit FGFR1 on the apical surface of Human Brain Microvascular Endothelial Cells (HBMECs)

To study the possible interaction of FGFR1 by meningococal colonies, HBMECs were infected with *N. meningitidis* for 4 h, fixed, and prepared for immunofluoresce microscopy. FGFR1 was recruited by meningococcal colonies on the apical surface of HBMECs. FGFR1 coincided with recruitment of both 67LR and 37LRP isoforms of laminin receptor (Figure 1). To address whether FGFR1 recruited by meningococci is activated, HBMECs were labelled with primary antibodies specific for phosphorylated tyrosine 766 (p-Y766), 67LR and 37LRP. In all cases the phosphorylated FGFR1 co-localised with meningococcal cells (Figure 1).

**Figure 1.**
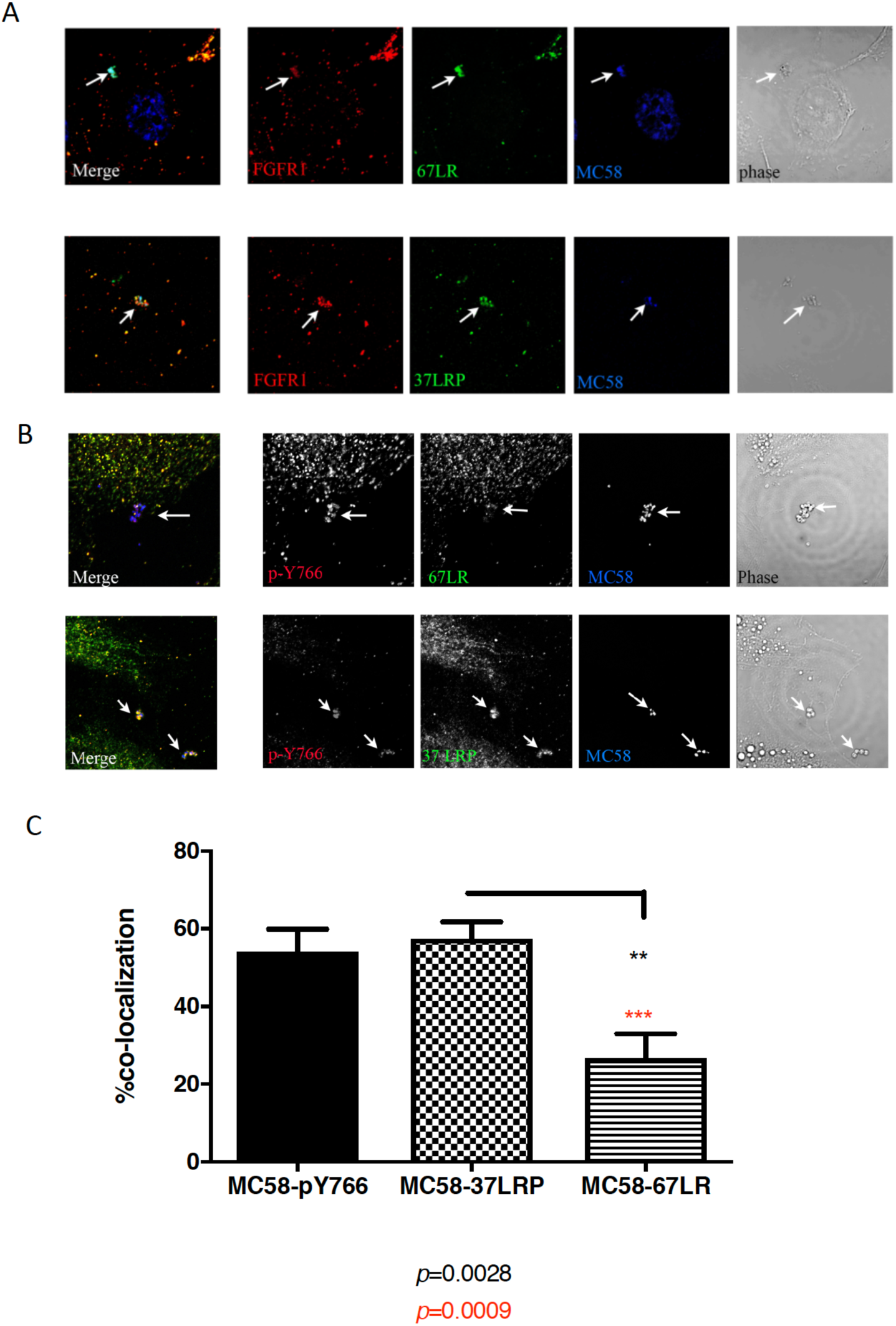
Activated FGFR1 is recruited by *N. meningitidis* colonies. (A) HBMECs were infected with meningococcal cells (MOI: 200) for 2-4 h. *N. meningitidis* colonies were visualized with DAPI (blue), FGFR1 was probed with anti-FGFR1 primary antibody and detected with anti-goat Alexa Fluor 680 antibody (red). 37LRP and 67LR were probed with primary antibodies and detected with anti-mouse Alexa Fluor 488 antibodies (green). FGFR1 was recruited by meningococcal colonies, which coincides with recruitment of 37LRP and, to a less extent, 67LR. Co-localisation area is shown by arrows (images are representative of 10 infected cells). (B) FGFR1 phosphorylated at Tyrosine 766 (p-Y766) was labelled with Alexa Fluor 680 (red) and both isoforms of Laminin receptor (67LR and 37LRP) with Alexa Fluor 488 (green). Recruitment of activated FGFR1 (p-Y766) coincided with recruitment of 37LRP and, to a less extent, 67LR. Levels of co-localization of MC58 with 37LRP, 67LR and p-Y766 (activated FGFR1) were quantified by measuring the percentage of co-localisation of each receptor with MC58 in 30 fields. There was a significant difference between recruitment of activated FGFR1 and 37LRP with recruitment of 67LR (*p=* 0.0009 and *p=* 0.0028 respectively; two tailed unpaired t-test).

Meningococcal colonies co-localised with activated FGFR1 and 37LRP, up to 60% of which was co-localised with FGFR1. This was significantly higher than co-localisation of the microcolonies with the 67LR isoform of the receptor (Figure 1-B, C).

### Internalised N. meningitidis associated with activated FGFR1

To determine whether internalised bacteria are associated with activated FGFR1, HBMECs were infected for 4 h with *N. meningitidis* and non-internalised bacterial cells were killed by gentamicin. Immunofluorescent staining for actin and activated FGFR1 (p-Y766) confirmed that meningococci recruited activated FGFR1 in the cytoplasm of HBMECs, and that the receptor was trafficked inside the cells alongside with meningococcal cells (Figure 2). To confirm that the bacterial cells in gentamycin-treated monolayers were internalised, a Z-stack image was constructed. Bacterial cells could be observed beneath the membrane of the endothelial cells (Figure 2-B). Furthermore, when cells were permeabilised prior to gentamycin treatment no bacterial cells were observed (Figure 2-C).

**Figure 2.**
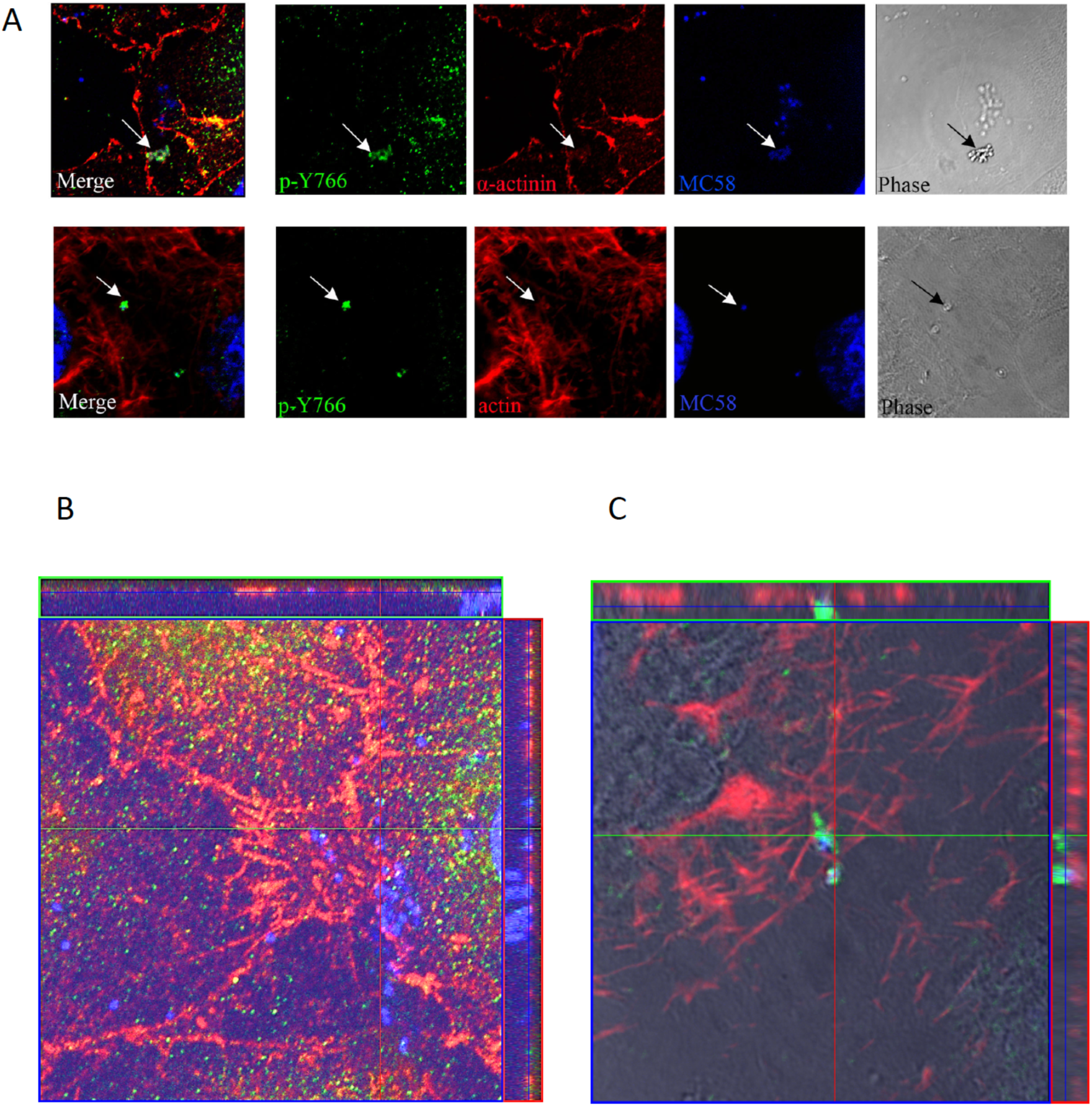
Activated FGFR1 is recruited by internalised meningococci within HBMECs. (A) In infected HBMECs α-actinin and actin were labelled with Alexa Fluor 680 (Red) and the activated form of FGFR1 (p-Y766) labelled with Alexa Fluor 488 (Green). MC58 was labelled with DAPI (Blue). Internalised bacteria co-localised with both α-actinin and activated FGFR1 (p-Y766) (arrow heads). (B and C) Z-stack image of meningococcal colonies showes that internalised bacteria (co-localising with α-actinin) recruit activated FGFR1 (p-Y766) inside the cells.

### FGFR1 expression and activation is required for meningococcal invasion into HBMECs

To study the role of FGFR1 in interaction of meningococci with HBMECs, FGFR1 expression was knocked down in HBMECs using siRNA treatment. Sixty hours post-siRNA treatment, cells were infected with meningococci and cell association and invasion assays were performed. The chemical inhibitor of FGFR1 (SU5402) (Mohammadi *et al.*, 1997) was also used to examine the effect of inhibiting the activation of FGFR1 during meningococcal infection.

The numbers of meningococcal cells associated with HBMECs was significantly and dramatically reduced in response to either FGFR1 knock-down or SU5402-treatment (Figure 3-A). Treatment with scrambled siRNA under the same conditions did not result in a significant reduction in association of meningococci with HBMEC cells. There was also a significant decrease in the number of internalised meningococcal cells recovered from FGFR1 siRNA-transfected HBMECs, as well as SU5402-treated HBMECs (Figure 3-B). Again, treatment with a scrambled siRNA did not significantly affect binding of meningococci to the HBMEC cells. This demonstrates that FGFR1 expression and activation plays an important and specific role in meningococcal adhesion to and invasion into HBMECs (Supporting material, S3).

**Figure 3.**
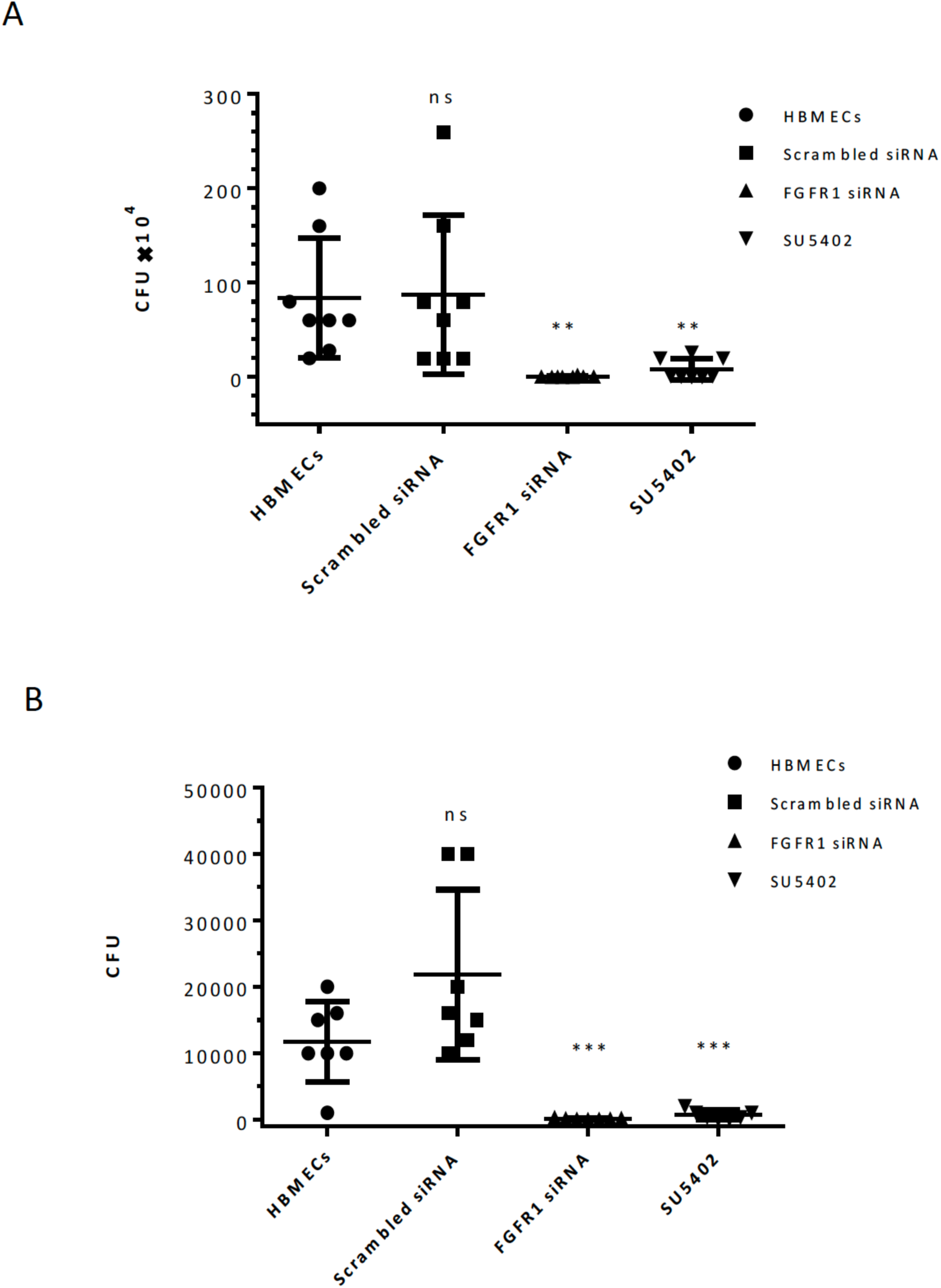
Expression and activation of FGFR1 plays an important role in the invasion of meningococci into HBMECs. (A) HBMECs were infected with *N. meningitidis* MC58 for 4 h (MOI: 10). Cells were washed and lysed in 500 µl of 1% saponin in PBS and 100 µl of homogenized lysates used for serial dilution preparation (up to 10 fold). 10 µl of each dilution was plated onto chocolate agar and CFUs were calculated for each sample. There was a significant reduction in the number of meningococci associated with HMBECs after FGFR1 knockdown (FGFR1 siRNA; *p*=0.0031, two tailed unpaired t-test, n=8) or in SU5402-treated cells in which activation of FGFR1 is chemically inhibited (*p*=0.0054, two tailed unpaired t-test, n=8; experiments were performed in triplicate wells and means shown represent 8 independent experiments). (B) For invasion assays, gentamicin was added after 4 h of infection and plates were incubated for a further 1 h. Cells were then washed, lysed, homogenised and dilutions plated onto chocolate blood agar plates. There was a significant difference between the number of internalised meningococci in FGFR1 siRNA-transfected cells compared to untreated cells, or cells treated with scrambled siRNA (*p*=0.0003 two tailed unpaired t-test, n=7). Chemical inhibition of FGFR1 activation also inhibited internalization of meningococcal cells into HBMECs (*p*=0.0005 two-tailed unpaired t-test, n=7, the error bars represent standard deviation of mean).

### There is a direct and specific interaction between FGFR1 IIIc and N. meningitides

To determine which isoforms of FGFR1 are expressed in HBMEC cells we performed RT-PCR using cDNA generated from total RNA extracted from HBMECs. FGFR1 IIIc and FGFR3 IIIb isoforms were both expressed in these cells (Supporting material, S1). To determine whether meningococci could interact directly with the extracellular domain of FGFR1 IIIc this protein was cloned and expressed as an Fc-tagged fusion protein (Supporting material). Two other proteins: Fc-FGFR2 IIIa TM, comprising the trans-membrane region of FGFR2 IIIa fused to the immunoglobulin Fc domain, and the Fc portion of immunoglobulin alone (Fc-stop) were used as controls for possible interaction with *N. meningitidis* that was not specific to the FGFR1 IIIc (extracellular) domain. Both control proteins were derived from clones employing the same vector and purified by the same method as Fc-FGFR1 IIIc. The purified proteins were employed as an immobilised ligand in ELISA experiments.

ELISA plates were coated with Fc-tagged purified proteins then binding of DIG-labelled *N. meningitidis* (MC58) was assessed (Figure 4, A and B). *N. meningitidis* (MC58) bound Fc-FGFR1 to a significantly greater degree than either Fc-FGFR2 IIIa TM or Fc-stop (Figure 4 A). This indicates that the observed interaction of Fc-FGFR1 was not due to meningococci binding to the Fc-tag of the expressed extracellular domain of FGFR1 and thus indicates a direct interaction between the receptor present on the apical surface of HBMECs and surface structures of *N. meningitidis.*

**Figure 4.**
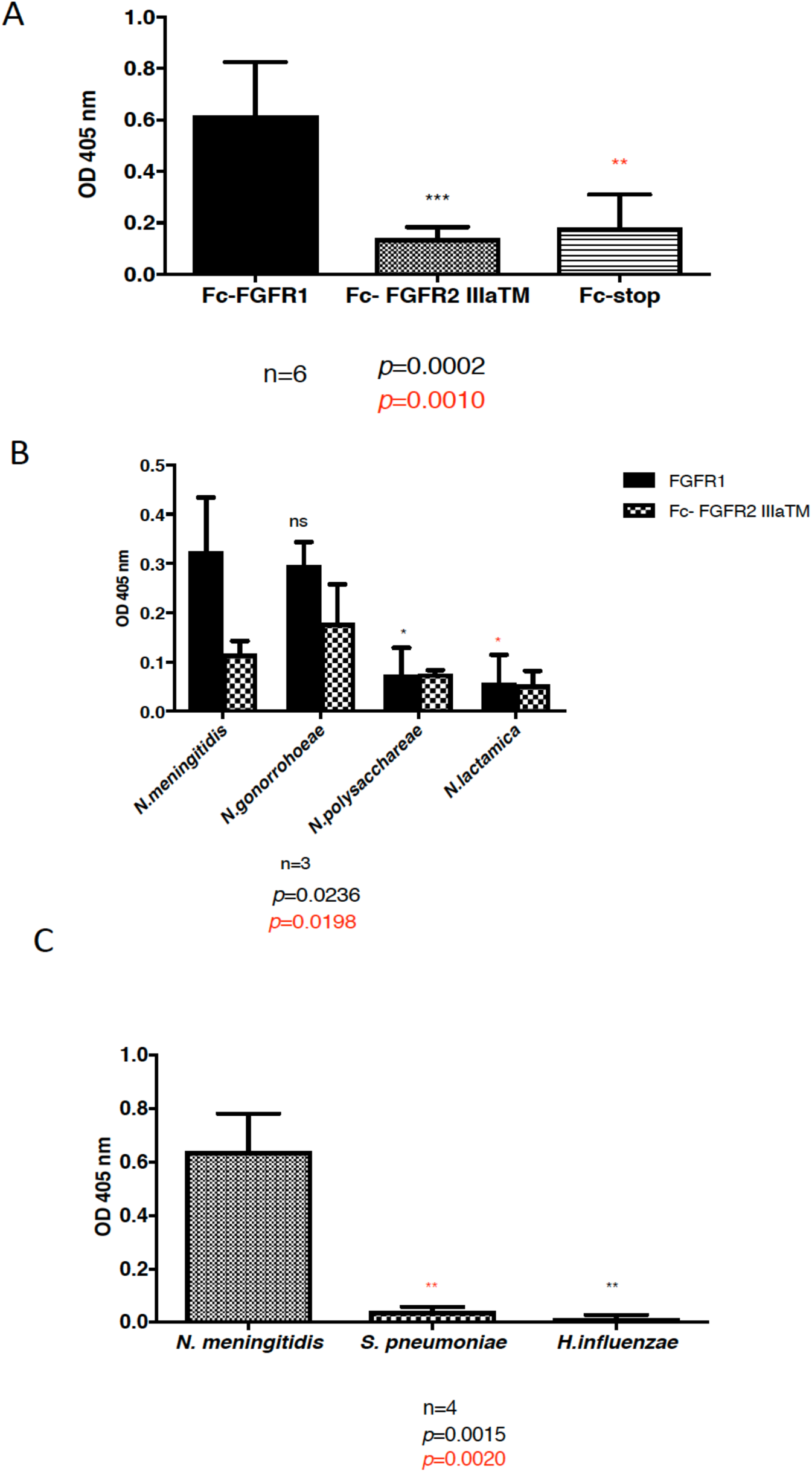
*N. meningitides* and *N. gonorrhoeae* interact directly with the extracellular domain of FGFR1. Fc-tagged purified proteins were used as the immobilised ligand in ELISA experiments. Levels of interaction with DIG-labelled MC58 were measured; the values shown are those following subtraction of binding to 1% BSA/PBS. (A) *N. meningitidis* MC58 interacts directly with the extracellular domain of Fc-FGFR1 (two tailed t-test; each experiment was performed in 6 technical replicates and the data shown is derived from 6 independent experiments). (B) Binding of commensal *Neisseria,* but not *N. gonorrhoeae,* to FGFR1 IIIc was significantly lower than that shown by *N. meningitidis* MC58. (C) The meningeal pathogens *H. influenzae* and *S. pneumoniae* bound FGFR1 IIIc to a negligible degree that was significantly lower than cells of *N. meningitidis* MC58 (*p*<0.002).

### Interaction between N. meningitidis and FGFR1 is also seen in the pathogen N. gonorrhoea but not commensal Neisseria species, nor other bacterial pathogens targeting the meninges

Having shown that Fc-FGFR1 IIIc interacts with meningococci, we sought to establish whether other *Neisseria* species could interact with this receptor. Representative strains belonging to several *Neisseria* species including the normally commensal species *N. polysaccharea* and *N. lactamica*, and the pathogenic *N. gonorrhoeae* were tested for interaction with Fc-FGFR1 in ELISA assays. *N. gonorrhoeae* cells bound to wells containing Fc-FGFR1 at similar levels to *N. meningitidis* MC58, and in both cases to a significantly higher degree than either of the two commensal species, which did not bind to a significantly higher degree to Fc-FGFR1 than to control wells containing only Fc-FGFR2 TM (Figure 4 B). We previously showed that, like the meningococcus, the meningeal pathogens *S. pneumoniae* and *H. influenzae* each targeted the laminin receptor on HBMEC cells as a prerequisite for internalisation (Orihuela *et al.*, 2009). We examined the possible interaction between these two other major causes of bacterial meningitis and the extracellular domain of FGFR1. Neither representative *S. pneumoniae* nor *H. influenzae* strains bound significantly to Fc-FGFR1 (Figure 4 C), demonstrating that the observed interaction between the pathogenic *Neisseria* species was specific.

## Discussion

The requirement for Focal Adhesion Kinases and activation of Src in the internalization of meningococci via interaction with integrins has been reported previously (Slanina *et al.*, 2012). Considering the role of FGFR1 in maintaining the integrity of the BBB and angiogenesis (van Hinsbergh *et al.*, 2005), we examined the possible role of FGFR1 in interactions with meningococci. Confocal microscopy studies on cells infected with meningococci confirmed that these bacteria recruit FGFR1 to the apical surface of HBMECs. Recently, we showed that meningococci bind to both 37LRP and Galectin-3 on the surface of HBMECs (Alqahtani *et al.*, 2014). Here, we showed that FGFR1 recruitment coincided with recruitment of both isoforms (37LRP and 67LR) of the laminin receptor; molecules already implicated in Neisserial-HBMEC interactions (Orihuela *et al.*, 2009). Ligation of the extracellular domain of FGFRs by their ligands leads to auto-phosphorylation of tyrosine residues in the cytoplasmic domain of the receptor; these phosphorylated residues subsequently serve as docking sites for a number of adaptor proteins responsible for regulation of various downstream signalling cascades (Turner *et al.*, 2010). The FGFR1 molecules recruited by meningococci were shown to be activated and activated receptors and meningococcal cells also co-localised with α-actinin. This is in agreement with previous studies on the trafficking of FGFR1 into early endosomes inside the cytoplasm, and the regulation of its trafficking by Syndecan 4 in a clathrin-independent manner (Elfenbein *et al.*, 2012). We also showed that the whole receptor is internalised into the cytoplasm along with invading meningococci; this was also observed by confocal microscopy in HBMECs treated with gentamicin after 4 h of meningococcal infection. Interestingly, FGFR1 engagement by basic fibroblast growth factor receptor has recently been shown to protect the integrity of HBMEC monolayers preventing the downregulation of the junction proteins zO-1, occludin and VE-cadherin in response to oxygen-glucose deprivation and deoxygenation (Lin *et al.*, 2017). This might have implications for the route of entry of meningococci via a trans-cellular pathway through HBMECs rather than a para-cellular pathway in which the integrity of the monolayer would have to be compromised.

To determine whether FGFR1 was required for meningococcal-HBMECs interactions, FGFR1 expression was transiently inhibited by using FGFR1 siRNA transfection and in other experiments its activity was inhibited by the specific chemical inhibitor SU5402. Both FGFR1 knock-down and SU5402 treatment of HBMECs resulted in a dramatic reduction in both association and internalisation of meningococci into HBMECs. Our data confirm that direct interaction between the extracellular domain of FGFR1 and meningococci is required for consequent activation of the receptor and internalisation of bacteria into the HBMECs.

The mechanisms by which recruitment of FGFR1 by meningococcal colonies leads to their internalisation is unknown. Interaction of meningococci with HBMECs has previously been shown to lead to higher levels of activation of ERK 1, 2 due to activation of ErbB2 in these cells (Hoffmann *et al.*, 2001). However, levels of ERK 1,2 activation in cells in which FGFR1 was knocked down by siRNA transfection were unaffected, demonstrating that FGFR1 does not regulate the levels of ERK1, 2 activation (Estes *et al.*, 2006).Several studies on meningococcal infection of endothelial cells showed that invasion of bacteria requires activation of Src, phosphorylation of cortactin via the Src pathway and activation of focal adhesion kinases (FAKs) (Hoffmann *et al.*, 2001, Miller *et al.*, 2012, Slanina *et al.*, 2012). Also, it has been shown that meningococcal cells hijack the β-arrestin/β 2-adrenoreceptor pathway to invade endothelial cells and cross the BBB: inhibition of β-arrestin mediated activation of Src, prevents the invasion of meningococcal cells (Coureuil *et al.*, 2010). Src is required for cortactin phosphorylation by FGF1 which can provide an alternate downstream pathway of FGFR1 from PLCγ and can be involved in cytoskeletal rearrangement (Zhan *et al.*, 1994, Liu *et al.*, 1999). However it was reported that mutation of Y766 in FGFR1 leads to higher level activations of PLCγ which inhibits Src activation (Landgren *et al.*, 1995). These observations suggest that FGFR1 siRNA transfection of HBMECs and chemical inhibition of FGFR1 (SU5402 treatment) may have led to the same effect on inhibition of Src activation which consequently inhibited meningococcal invasion into HBMECs. It is likely that FGFR1 plays an important role in meningococcal interaction with the BBB during infection. This effect appears to be specific to the meningococcus as the bacterial pathogens *H. influenzae* and *S. pneumoniae*, which also cross the BBB and can cause meningitis, do not interact with FGFR1 IIIc on the surface of endothelial cells. On the other hand, *N. gonorrhoea*, which does not usually interact with the BBB is able to bind this receptor. The significance of this is unknown. Further investigations are required to understand the role of FGFR1 signalling in meningococcal invasion into HBMECs.

## Experimental Procedures

### Bacterial growth and culture

*N. meningitidis* serogroup B strain MC58 was obtained from the American Type Culture Collection (ATCC) (Tettelin *et al.*, 2000) and routinely cultured on chocolated horse blood agar (Chocolate agar; Oxoid). *H. influenzae* Rd KW20 (ATCC 51097) (Fleischmann *et al.*, 1995) and *S. pneumoniae* T4R (unencapsulated) (Fernebro *et al.*, 2004) were also cultured on chocolated horse blood agar (Oxoid). All three bacteria were grown at 37°C, in an atmosphere of air plus 5% CO_2_.

### Cell association assay and cell invasion (Gentamicin protection) assay

To quantify cell association and cell invasion of HBMECs with *N. meningitidis* HBMECs were seeded and grown overnight or for 48 h after siRNA transfection until 100% confluent in 24-well plates. Cells were infected with 1×10^7^ CFU bacteria for 4 h in ECM-b media without any supplements. Cell association and invasion were then determined as described previously (Oldfield *et al.*, 2007).

### Confocal Immunofluorescent Microscopy

HBMECs were seeded onto fibronectin-coated coverslips (1-10 × 10^5^ cells) and grown overnight to reach a confluency of 70-80%. Cells were infected for 2-4 h (MOI 200-300). Coverslips were washed with PBS and fixed with 4% paraformaldehyde (w/v) in PBS for 5 min. Coverslips were then washed with PBS and blocked in 4% (w/v) BSA/PBS at 4°C overnight. For intracellular staining, cells were permeabilised by treatment with 0.1% Saponin, 20 mM glycine in 4% BSA/TBS at 4°C. Subsequent staining procedures were carried out in r 4% (w/v)BSA/TBS. Briefly, coverslips incubated with primary antibody for 1 h were washed with PBS followed by one wash with dH_2_O. Coverslips were then incubated with secondary antibody for 1 h in the dark followed by washes with PBS-Tween (0.05% v/v; PBS-T), PBS and then dH_2_O. Coverslips were then mounted on glass slides with ProLong® Gold and SlowFade® Gold Antifade Reagents with DAPI (Invitrogen). Coverslips were analysed using a Zeiss LSM-700 confocal microscope. Images were processed with ImageJ, Adobe Photoshop and LSM Image Browser software.

### Antibodies and reagents

Antibodies detecting FGFR1 (Flg S-16 and Flg C-15), phosphorylated FGFR1 (p-Y766) were purchased from Santa Cruz Biotechnology. Secondary antibodies conjugated to various fluorochromes, and Phalloidin conjugated to fluorochrome 488 were obtained from Life Technologies-Invitrogen.

Antibody detecting 37LRP (A-7) was purchased from Santa Cruz Biotechnology and antibody against 67LR (Mluc-5) was purchased from Thermo Scientific. FGFR1-specific inhibitor SU5402 (Mohammadi *et al.*, 1997) was purchased from Calbiochem.

Fc-tagged FGFR1 IIIc (*ca.* 80 kDa), Fc-tagged FGFR2 IIIa TM (*ca.* 61.9 kDa) and Fc-tag (*ca.* 29.26 kDa) were expressed and purified using a protein A column (S1).

### FGFR1 siRNA transfection in HBMECs

Human FGFR1 siRNA (siGENOME SMART pool) and control scrambled siRNA were obtained from Dharmachon/Thermo Scientific and reconstituted following the manufacturer guidance. FGFR1 siRNA was resuspended in 1 ml of 1 × siRNA buffer to a final concentration of 5 µM (stock). HBMECs were seeded into 24-well plates pre-coated with fibronectin, as previously described, and grown overnight to reach a confluency of 70-80%. Transfection media was prepared by mixing serum- and antibiotic-free media with siRNA from a 50µM stock to a final concentration of 5µM; in a separate tube Transfection reagent number 1 (Thermo Scientific-Dharmacon) was added to serum and antibiotic-free media. Both tubes were incubated for 5 min at room temperature and then mixed together by pipetting and incubated at room temperature for 20-30 min. Cells were washed with serum and antibiotic-free medium and 240 µl of complete media without antibiotics were added to each well and then the transfection mixture was added drop-wise to each well to a final concentration of 50 nM siRNA/ well. Cells were then incubated for 6 h and then the media was replaced with complete media (Endothelial cell medium (ECM-b) (ScienCell) supplemented with ECGS (containing EGF, VEGF) (ScienCell) (1% v/v) and FBS (5% v/v) and penicillin/streptomycin (ScienCell; 1% v/v). The level of FGFR1 expression was examined at RNA and protein levels at 24, 48 and 72 h post transfection.

### FGFR1 inhibition in HBMECs

HBMECs were serum-starved and treated with SU5402 (0.5 µM) 1 h prior to infection. Approximately 10×10^6^ / well (MOI: 10) of bacteria were added to each well and infected cells were incubated for 4 h at 37°C/5% CO2 in medium containing 0.5 µM SU5402. Cells were then washed twice with PBS and lysed in 1% saponin/PBS. Cell lysates were homogenised and appropriate dilutions plated out to calculate the levels of association. For invasion assays, cells were treated with gentamicin for 1 h to kill non-internalised bacteria.

### ELISA

96-well plates (NUNC Immobilizer Amino) were coated with 100 µl of protein A (Pierce; 1 µg ml^−1^) in PBS for 1 h, washed once with PBS-T and then 86.5 ×10 ^−15^ M of Fc-tagged recombinant proteins added to each well in carbonate buffer (pH 9.6). Plates were incubated for 1 h and then washed three times with PBS-T. Plates were blocked with 1% BSA/PBS (w/v) for 1 h, then washed once with PBS-T. Bacterial cells harvested from overnight plates and resuspended in PBS-T, washed with the same buffer three times and finally resuspended in sodium carbonate buffer (44 mM NaHCO_3_, 6.0 mM Na_2_CO_3_; pH: 9.6). The OD_600_ was measured and 20 ng (2 µl of 10 ng µl^−1^) of digoxigenin (DIG; Roche) was added to 1 ml of bacterial suspension with OD_600_:1. The bacterial suspensions were incubated for 2 h in the dark at room temperature on the shaker. Bacteria then were washed three times with PBS-T by centrifugation (13000 × *g* for 1 minute) and resuspended in 1% BSA/PBS (w/v). OD_600_ in 1% BSA/PBS was adjusted to 0.02 for ELISA. For each experiment fresh labelled bacterial strains were used. 100 µl of DIG-labelled bacteria were added to each well and plates were incubated at 4°C overnight then washed five times with PBS-T. 100 µl of anti-DIG-alkaline phosphatase antibody (Roche; 0.0002 v/v) in 1% BSA/PBS was then added to each well and incubated for 1 h then washed three times with PBS-T. 200 µl of alkaline phosphatase substrate (SIGMA) was added and plates were incubated for 1 h. The OD_405_ was measured for each sample and values obtained subtracted from the binding of the same DIG-labelled strain to 1% BSA/PBS.

## Acknowledgements

We thank Dr S. Morroll for constructing the plasmid encoding Fc-Stop-LRP.

## Funding Statement

This work was funded by the Medical Research Council, UK (www.mrc.ac.uk); award number G0801173. The funders had no role in study design, data collection and analysis, decision to publish, or preparation of the manuscript.

## Supporting material

### HBMECs express FGFR1 IIIc and FGFR3 IIIb isoforms

HBMECs were grown overnight, RNA extracted and cDNA prepared. This was then used as template to amplify the third IgG-like domain of FGFRs (exon 7, 8 IIIb) and (exon 7, 9 IIIc) isoforms (Hajihosseini *et al.*, 1999). RT-PCR showed that HBMECs express FGFR1 IIIc and FGFR3 IIIb isoforms.

**S1.**
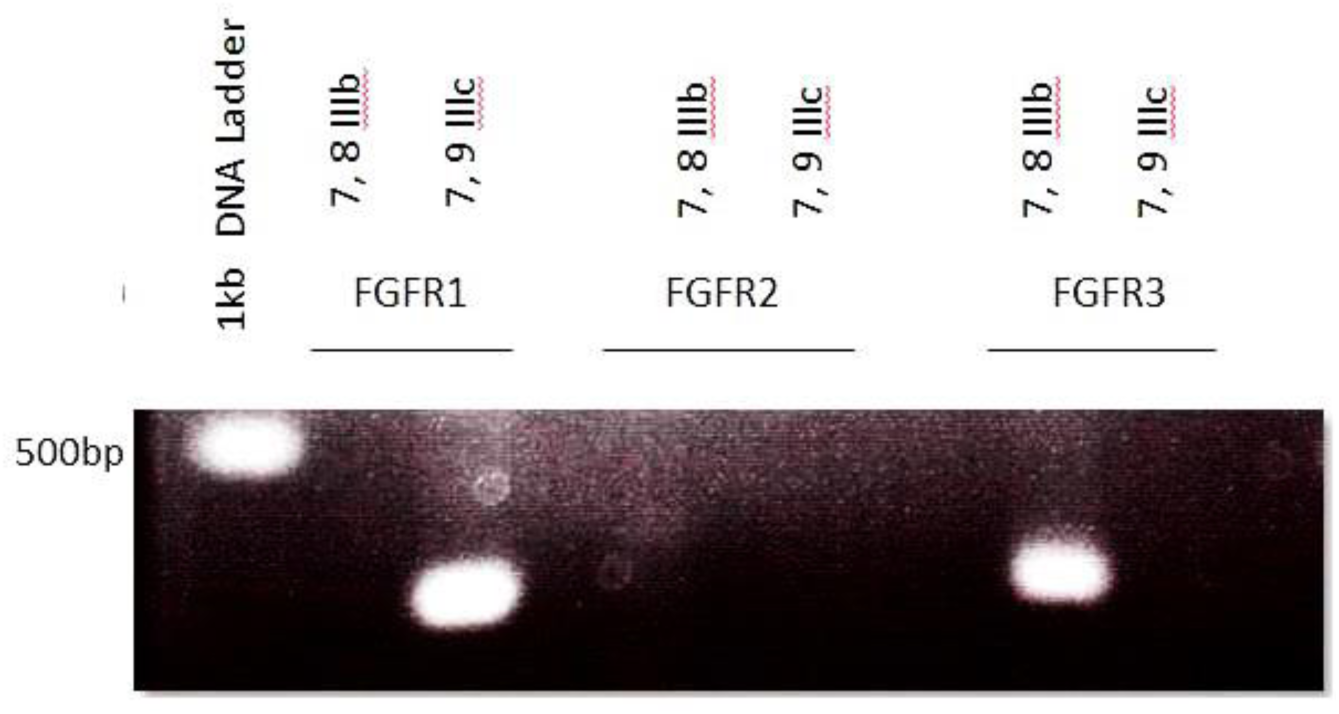
HBMECs express FGFR1 IIIc and FGFR3 IIIb isoforms. RT-PCR performed using cDNA generated from total RNA extracted from HBMECs. 100ng of cDNA was used in as template in each PCR reaction. Various combinations of primers specific for exon 7, 8 and 9 were used to determine the type of FGFR expressed by HBMECs.

### Construction of FGFR1 IIIc into pEF-Bos-ss-Fc-ires-TPZ mammalian vector

HBMECs were routinely grown in fibronectin-coated T75 flasks (BD Bioscience). For RNA extraction, cells were harvested by trypsin treatment. Cell suspensions were then centrifuged for 5 min at 300 × *g*, the supernatant was discarded and cells were lysed in lysis buffer (SIGMA GenElute^™^ Mammalian Total RNA Miniprep Kits) supplemented with 1% (v/v) 2-β-Mercaptoethanol and RNA extracted according to the manufacturer’s protocol (Sigma GenElute^™^ Mammalian Total RNA Miniprep Kit). The concentration of extracted RNA was measured using a Nanodrop spectrophotometer and adjusted to 200 ng µl^−1^. To remove traces of genomic DNA from RNA samples, DNAse treatment was performed following the manufacturers protocol (Turbo DNase, Life Technologies). 10 µl of RNA was then used for cDNA preparation using the High Capacity cDNA Reverse Transcription kit (Applied Biosystem). HBMEC cDNA was then used as template to amplify the extracellular domain of FGFR1 IIIc isoform. Primers were designed to amplify the extracellular domain of FGFR1 containing restriction sites for NdeI and NotI enzymes. The amplified PCR product was gel purified, digested and then ligated into NotI and NdeI-digested pEF-Bos-ss-Fc-FGFR2IIIaTM-ires-TPZ (Mizushima et al., 1990, Burgar et al., 2002, Wheldon et al., 2011) using the LigaFast TMRapid DNA Ligation System (Promega). The ligated plasmid was used for transformation of competent *E. coli* JM109 cells (Promega). Plasmids extracted from ampicillin-resistant transformants were screened by restriction digestion and DNA sequencing. The plasmid from one correct clone was named pEF-Bos-ss-Fc-extFGFR1IIIc-ires-TPZ.

**S2.**
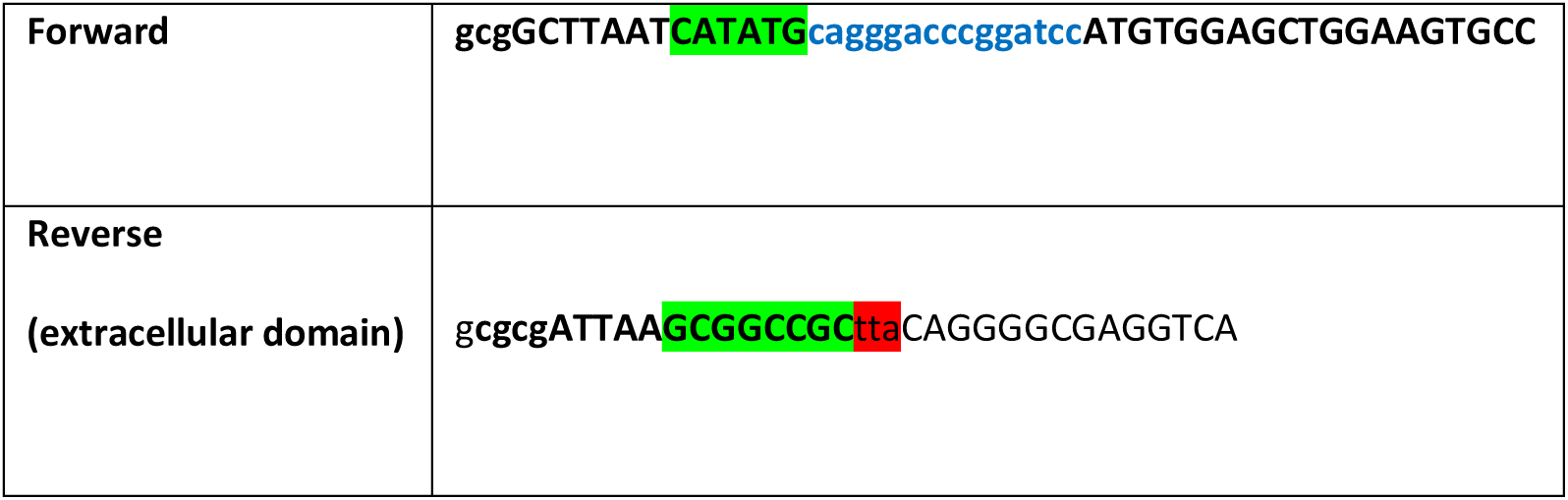
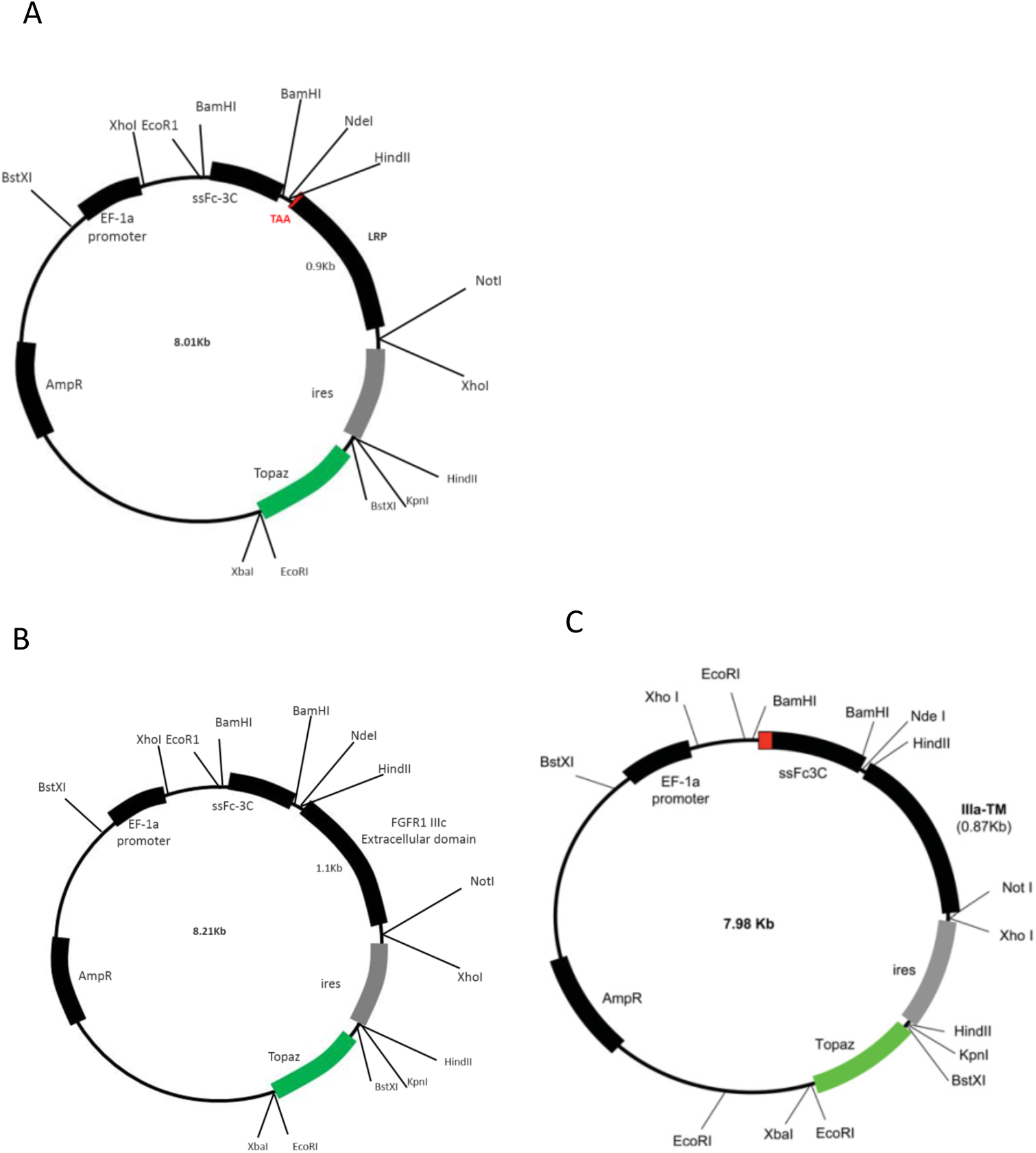
Schematic view of the plasmid pEF-Bos-ss-Fc-extFGFR1IIIc-ires-TPZ. The extracellular domain of FGFR1 IIIc (A) and LRP were replaced with FGFR2 IIIaTM insert of the plasmid pEF-Bos-ss-Fc-FGFR2 IIIa TM-ires-TPZ (C) (Wheldon *et al.*, 2011). Stop codon (TAA in Red-B) was introduced in start of the LRP coding gene to be able express the Fc-tag and not the LRP protein (B). The plasmid encoding Fc-Stop-LRP was constructed in house by Dr. S. Morroll.

### Purification of Fc-tagged recombinant proteins

Media was collected 72 h post-transfection of 293T cells and Fc-tagged recombinant proteins purified on a Protein AIG-Sepharose (Source BioScience LifeSciences) column. Briefly, the media was diluted 1:1 (v/v) with binding buffer (0.05 M sodium borate, 0.15 M sodium chloride; pH: 8). 100 µl of beads were added to 30ml of diluted media and incubated with shaking at 4°C overnight. The column was prepared and washed 4-5 × with binding buffer. The column was then washed with binding buffer containing 0.5 M sodium chloride. Proteins were eluted in 1 ml of 0.1 M glycine (pH: 2.5) and neutralised with 100 µl of 1 M Tris-HCl (pH: 9) per 1 ml of eluted samples. A total number of 9-10 protein fractions were eluted and stored at −80°C. Protein concentration in each fraction was quantified using the BCA kit following manufacturer’s protocol (Thermo Scientific).

**S3.**
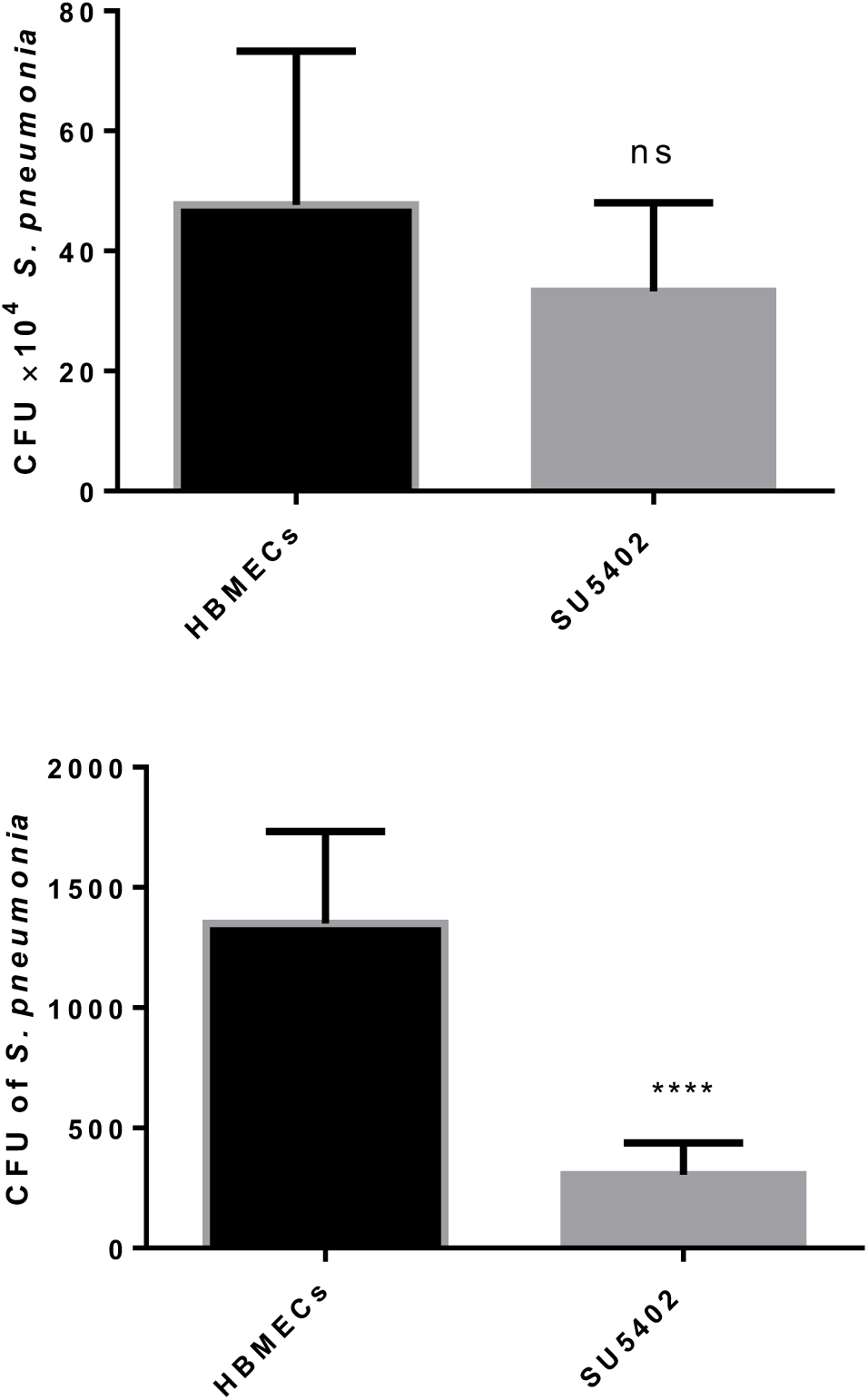
Chemical inhibition of FGFR1 does not affect the number of *S. pneumoniae* associated with HBMECs but reduced the number of invaded bacteria into HBMECs. HBMECs were infected with *S. pneumonia* at MOI: 10. Cell association and invasion assays were performed and the CFUs were calculated.

